# POPULATION GENOMIC ANALYSIS OF *CRYPTOCOCCUS* BRAZILIAN ISOLATES REVEALS AN AFRICAN TYPE SUBCLADE DISTRIBUTION

**DOI:** 10.1101/2021.02.08.430197

**Authors:** Corinne Maufrais, Luciana de Oliveira, Rafael W. Bastos, Frédérique Moyrand, Flavia C. G. Reis, Clara Valero, Bianca Gimenez, Luisa J. Josefowicz, Gustavo H. Goldman, Marcio L. Rodrigues, Guilhem Janbon

**Author notes:** Both authors should be considered as first authors (Corinne Maufrais, Luciana de Oliveira).

## Abstract

The genomes of a large number of *Cryptococcus neoformans* isolates have been sequenced and analyzed in recent years. These genomes have been used to understand the global population structure of this opportunistic pathogen. However, only a small number of South American isolates have been considered in these studies, and the population structure of *C. neoformans* in this part of the world remains elusive. Here, we analyzed the genomic sequences of 53 Brazilian *Cryptococcu*s isolates and deciphered the *C. neoformans* population structure in this country. Our data reveal an African-like structure that suggested repeated intercontinental transports from Africa to South America. We also identified a mutator phenotype in one VNBII Brazilian isolate, exemplifying how fast-evolving isolates can shape the *Cryptococcus* population structure. Finally, phenotypic analyses revealed wide diversity but not lineage specificity in the expression of classical virulence traits within the set of isolates.

---

The current classification identifies seven human-pathogenic *Cryptococcus* species, which are responsible for 180,000 deaths every year in the world (Hagen *et al*. 2015; Rajasingham *et al*. 2017). *C. neoformans* and *C. deneoformans* represent the *neoformans* group of species, whereas the *gattii* group of species comprises *C. gattii, C. deuterogattii, C. tetragattii, C. decagattii*, and *C. bacillisporus* (Hagen *et al*. 2015). *C. neoformans* is the most common species in clinical settings, representing more than 95% of the clinical isolates (Kwon-Chung *et al*. 2014; Janbon *et al*. 2019). This species affects mostly immunocompromised patients in whom it provokes meningoencephalitis. Other species, such as *C. gattii*, can be primary pathogens and cause pulmonary infections (Kwon-Chung *et al*. 2014). *C. neoformans* is thought to be acquired very early in life and can stay in a dormant state for years, probably in alveolar macrophages (Garcia-Hermoso *et al*. 1999; Alanio 2020). As soon as a defect in the cellular immunity appears, yeast cells can multiply, reach the central nervous system, and induce meningoencephalitis, which is invariably deadly if untreated.

*C. neoformans* strains are ubiquitous in the environment and have been isolated from all continents except Antarctica. Similarly, although isolates of the *gattii* group of species have been long considered to be geographically restricted to tropical and subtropical regions and typically associated with *Eucalyptus* trees, they have emerged as a primary pathogen in western Canada and the United States of America (Kidd *et al*. 2004). Both species can live in a broad range of ecological habitats and have been commonly isolated from soil, guano, decaying wood, fruits, and insects (Lin and Heitman 2006). Pigeons (*Columba livia*) are vectors for the worldwide dissemination of *Cryptococcus*. Indeed, dry pigeon guano seems to be an ideal environment for *C. neoformans* development and is commonly infected with this fungus. Nevertheless, pigeons are usually not harmed by *C. neoformans* because their body temperature is too high (42.5°C) for its growth and disease development (Johnston *et al*. 2016).

All *neoformans* and *gattii* species are haploid species with two possible mating type alleles (*MAT***a** and *MATα*) and a bipolar mating system. Efficient sexual reproduction has also been described in the laboratory. Nevertheless, *C. neoformans MAT***a** isolates are rare in clinical and environmental isolates with the exception of some places in Africa (Kwon-Chung and Bennett 1978; Litvintseva *et al*. 2003). In addition, it has been reported that 8% of the population of *C. neoformans* isolates are diploid (Lin *et al*. 2009). Diverse hybrids have also been isolated from environmental and clinical sources. The most common are AD hybrids, which are produced by hybridization of *C. neoformans* with *C. deneoformans* isolates. In addition, rare cases of *gattii*/*neoformans* hybrids have been reported (Bovers *et al*. 2008; Aminnejad *et al*. 2012). Nevertheless, population structure analysis suggested that the different *neoformans* and *gattii* populations are largely clonal with a limited degree of recombination (Litvintseva *et al*. 2005; Bui *et al*. 2008; Litvintseva and Mitchell 2012).

Historically, these analyses were based on serotyping, which was first replaced by PCR-based characterization (RAPD, AFLP, mating type analysis) of the isolates and then by sequencing of a defined set of genes by multilocus sequence typing (MLST) (Ruma *et al*. 1996; Boekhout *et al*. 2001; Matsumoto *et al*. 2007; Meyer *et al*. 2009a). Originally, five *neoformans* molecular subtypes were thus defined (VNI, VNII, VNIV, VNIII, and VNB) (Meyer *et al*. 2009b). The new species definition includes VNI, VNII, and VNB within *C. neoformans*, VNIV within *C. deneoformans*, and VNIII as *C. neoformans*/*C. deneoformans* hybrids. VNI is the most common *C. neoformans* molecular subtype and has a global distribution. The VNB subtype was discovered in Africa and has a genetically variable population with 10% *MAT***a** isolates, which are very rare in all other studied populations (Litvintseva *et al*. 2003). The VNB clade includes strains isolated from the endemic mopane tree (*Colophospermum mopane*), whereas strains isolated from pigeon excreta in the same area had the VNI molecular type, which is shared by most isolates globally. This observation supported the “out of Africa” hypothesis in which few of these isolates would have adapted their biology to multiply in pigeons to achieve global dispersion (Litvintseva *et al*. 2011; Litvintseva and Mitchell 2012).

More recently, the improvement of sequencing technologies and bioinformatics tools allowed analysis of whole genome sequences, revealing new details in the structure and global geographical distribution of the *Cryptococcus* populations together with the identification of additional clades and subclades (Farrer *et al*. 2019). Most of these studies focused on *C. neoformans*, with >1000 *C. neoformans* genome sequences analyzed in four studies (Desjardins *et al*. 2017; Rhodes *et al*. 2017b; Vanhove *et al*. 2017; Ashton *et al*. 2019; Ergin *et al*. 2019). Through these efforts, the VNB clade has been separated into the VNBI and VNBII subclades, which have different resistance to oxidative stress and a different ability to produce melanin (Desjardins *et al*. 2017; Vanhove *et al*. 2017). The VNI clade has been separated into a number of subclades with three phylogenetic groups representing the majority of the studied isolates (Rhodes *et al*. 2017b; Ashton *et al*. 2019). Surprisingly, only 11 haploid strains isolated in South America were included in these studies. This small number conflicts with the relevance of South America as a source of genetic diversity in the *gattii* isolates (Engelthaler *et al*. 2014). Up until now, the only analysis of *C. neoformans* population structure in South America has been an MLST analysis that concluded that subtype ST93 was the most prevalent among strains isolated from HIV patients in Southeastern Brazil with a low level of recombination (Ferreira-Paim *et al*. 2017). Similarly, few phenotypic analyses that include South American isolates have been published, and a recent analysis of virulence factor expression within Brazilian *Cryptococcus* clinical isolates does not include any sequencing or molecular typing analysis (De Sousa *et al*. 2020). In the present study, we analyzed the phenotypes and genome sequences of 53 Brazilian isolates, revealing very high phenotypic diversity within *Cryptococcus* isolates and a subclade distribution that suggests several waves of *Cryptococcus* introduction from Africa to South America.

## Material and Methods

### Isolates used in this study

Strains 2B, 2C, 2D, 4C, 7, 33A1, 54A1, 56A1, 68B1, 69A1, 81A1, 96A1, 106A1, 144A1, 151A1, 176A1, 3Pb3, 5Pb2, 19Pb4, 23Pb2, 32Bp1, and RR 2605 were isolated from Northern Brazil (Dal Pupo *et al*. 2019) and kindly donated by Dr. Halan Dal Pupo. Strains Cn201, Cn359, Cn894, Cn14HC, and Cn201 were obtained from the culture collection of the Central Hospital of the University of São Paulo. All other isolates were obtained from the Collection of Pathogenic Fungi available at Fiocruz. No genomic information was available for any of these isolates before the present study.

### Read alignment, variant detection, and ploidy analysis

Illumina reads were aligned to the *Cryptococcus* reference genomes using Minimap2 aligner v. 2.9 (Li 2018) with the “-ax sr” parameter. BAM files were sorted and indexed using SAMtools (Li *et al*. 2009) version 1.9. Picard version 2.8.1 (http://broadinstitute.github.io/picard) tools were used to identify duplicate reads and assign correct read groups to BAM files. SAMtools version 1.9 and Picard version 2.8.1 were then used to filter, sort, and convert SAM files and assign read groups and mark duplicate reads. Single-nucleotide polymorphisms (SNPs) and insertions/deletions (indels) were called using Genome Analysis Toolkit version 3.6 with ploidy=1 according to the GATK Best Practices. HaploScore parameters used to filter SNPs and indels included VariantFiltration, QD <2.0, LowQD, ReadPosRankSum<-8.0, LowRankSum, FS >60.0, HightFS, MQRankSum<-12.5, MQRankSum, MQ <40.0, LowMQ, and HaplotypeScore >13.0. Phyml version 20120412 (Guindon *et al*. 2010) with the GTR, NNIs substitution model was used to infer phylogenetic relationships between the isolates using the separate dataset of confident SNPs. To examine variations in ploidy across the genome, the sequencing depth at all positions was computed using SAMtools (Li *et al*. 2009), and then the average depth was computed for 1-kb windows across the genome.

### MAT locus determination

The mating type of Brazilian isolates was determined as previously published (Rhodes *et al*. 2017b). Briefly, Illumina reads were aligned to the mating-type locus sequences [AF542529.2 and AF542528.2] using BWA-MEM. Computed average depths at the *SXI* and *STE20* loci using SAMtools mpileup were used to determine the mating type.

### Population inference by fastSTRUCTURE

We used fastStucture (Raj *et al*. 2014) to identify admixture in the dataset from K=2 to K=4. After K*=*4, clusters with the highest number of isolates were split into sub-clusters, which likely reflects an issue due to the number of isolates within clusters rather than true biological significance.

### Drug susceptibility testing

The minimum inhibitory concentration (MIC) of fluconazole (FLC) (Sigma-Aldrich) was determined as described by M27-A3 (Institute 2017).

### Titan cell production

Titan cell (TC) production was evaluated as described by Dambuza and colleagues with some modifications (Dambuza *et al*. 2018). The isolates were cultivated in yeast peptone dextrose (YPD) agar (1% yeast extract, 2% bacto-peptone, 2% glucose, 2% bacto-agar) for 48h at 30°C. A single colony was transferred to 5 mL YNB without amino acids (Sigma Y1250) prepared according to the manufacturer’s instructions plus 2% glucose. After incubation overnight at 30°C and 200 rpm, 1×10^3^ cells/mL were inoculated into 5 mL 1xPBS (phosphate buffer solution) + 10% heat-inactivated (HI) fetal calf serum (FCS) and incubated for 72 h at 37°C, 5% CO_2_. We visualized the cells microscopically without any staining, and 50 cells were measured. TCs were considered body cells (without the capsule) with a diameter >10 μm. The experiment was repeated twice.

### Capsule staining

Fungal cells were cultivated in YPD agar for 24 h at room temperature. Single colonies were transferred by sterile loop to the wells of a 96-well plate, each containing 200 μL of Roswell Park Memorial Institute (RPMI) medium at a final cell density of 10^7^ yeast/mL. The plates were incubated for an additional 24 h at 37°C in 5% CO_2_ and centrifuged to separate the cells from the supernatants. The supernatants were stored for GXM analysis, and the cells were washed twice in PBS for further fixation in 4% paraformaldehyde for 1 h at room temperature. Yeast cells were then blocked (1% bovine serum albumin [BSA] in PBS for 1 h at 37°C) and incubated with the 18B7 anti-GXM monoclonal antibody (Casadevall *et al*. 1992) (mAb 18B7; 10 μg/mL for 1 h at 37°C), a kind gift from Dr. Arturo Casadevall (Johns Hopkins University). The cells were washed and incubated with an Alexa Fluor 488-conjugated secondary antibody (10 μg/mL for 1 h at 37°C; Invitrogen, USA). The cells were finally stained with calcofluor white (5 μg/mL for 1 h at 37°C; Invitrogen, USA). Alternatively, the cells were counterstained with India ink for capsular visualization. The cells were visualized on a Leica TCS SP5 confocal microscope or analyzed with a FACS Canto II flow cytometer. Data were processed with the FACSDiva software, version 6.1.3.

### Data availability

All sequence data from this study have been submitted to GenBank under the BioProject identification no. PRJNA702892.

## Results

### Description of the origin of the Brazilian isolates

We selected 53 *Cryptococcus* Brazilian isolates, of which 12 were classified as *C. gattii* and 41 as *C. neoformans*. More than 41% (n=22) were environmental isolates and 36% (n=19) were clinical isolates. The origin of the isolate was unknown for the remaining 23% (n=12). Twenty-one strains were isolated from pigeon excreta in Northern Brazil (Dal Pupo *et al*. 2019), 28 isolates were obtained from the culture collections available at Fiocruz, and 4 were acquired from the Central Hospital of the University of São Paulo (Table S1).

### Genome characterization

Whole genome sequencing was performed for each of the 53 Brazilian isolates considered in this study. Between 2.7 and 10.9 million 100-nt paired-end reads were obtained and aligned to eight *Cryptococcus* reference genomes using Minimap2 (Li 2018) (Table S1). Reference genomes were *C. neoformans* strain H99 (Janbon *et al*. 2014), *C. deneoformans* strain JEC21 (Loftus *et al*. 2005; Gonzalez-Hilarion *et al*. 2016), *C. deuterogattii* strain R265 (Yadav *et al*. 2018; Gröhs Ferrareze *et al*. 2021), *C. gattii WM276, C. bacillisporus* strains CA1280 and CA1873, *C. decagattiii* strain AFLP10, and *C. tetragattii* strain IND107 (Basenko *et al*. 2018). Of the 53 isolates, 40 were identified as *C. neoformans* with >80% of the reads aligning to the H99 genome (<40% of reads could be aligned to any other reference genome). Among the 13 remaining isolates, 8 were *C. deuterogattii* and 5 were *C. gattii* (Table S1). Of note, we found that isolate Cg366, which was previously identified as *C. gattii*, is actually a *C. neoformans* isolate. Isolates Cn201 and Cn894, which were previously considered to be *C. neoformans*, were identified as *C. deuterogattii* isolates.

In order to determine mating type, we compared the number of reads aligned to the *STE20a, STE20alpha, SXI1*, and *SXI2* genes of the corresponding species. These analyses revealed that all isolates were *MATα*, confirming the overwhelming dominance of this mating type in clinical and environmental isolates observed in most areas in the world (Desjardins *et al*. 2017). The analysis of the read alignment profiles provided insight into possible large duplications or aneuploidies. All *C. neoformans* isolates displayed an even alignment profile. This observation together with the absence of heterozygous SNPs in our analysis suggested that these isolates are either haploid or euploid homozygotes. However, we observed a duplication of the end of the chromosome 1 in three isolates (Cn160, Cn161, and Cn186)) (left side in Figure 1). The isolate Cn161 apparently had additional duplications within chromosome 8. Similar patterns were observed in the eight *C. deuterogattii* isolates, with two isolates (Cn201 and Cn894) displaying a duplication of a region located in chromosomes 8 and 13, respectively. In contrast, all five *C. gattii* isolates displayed an even alignment profile, suggesting that all are haploid (data not shown).

**Figure 1.**
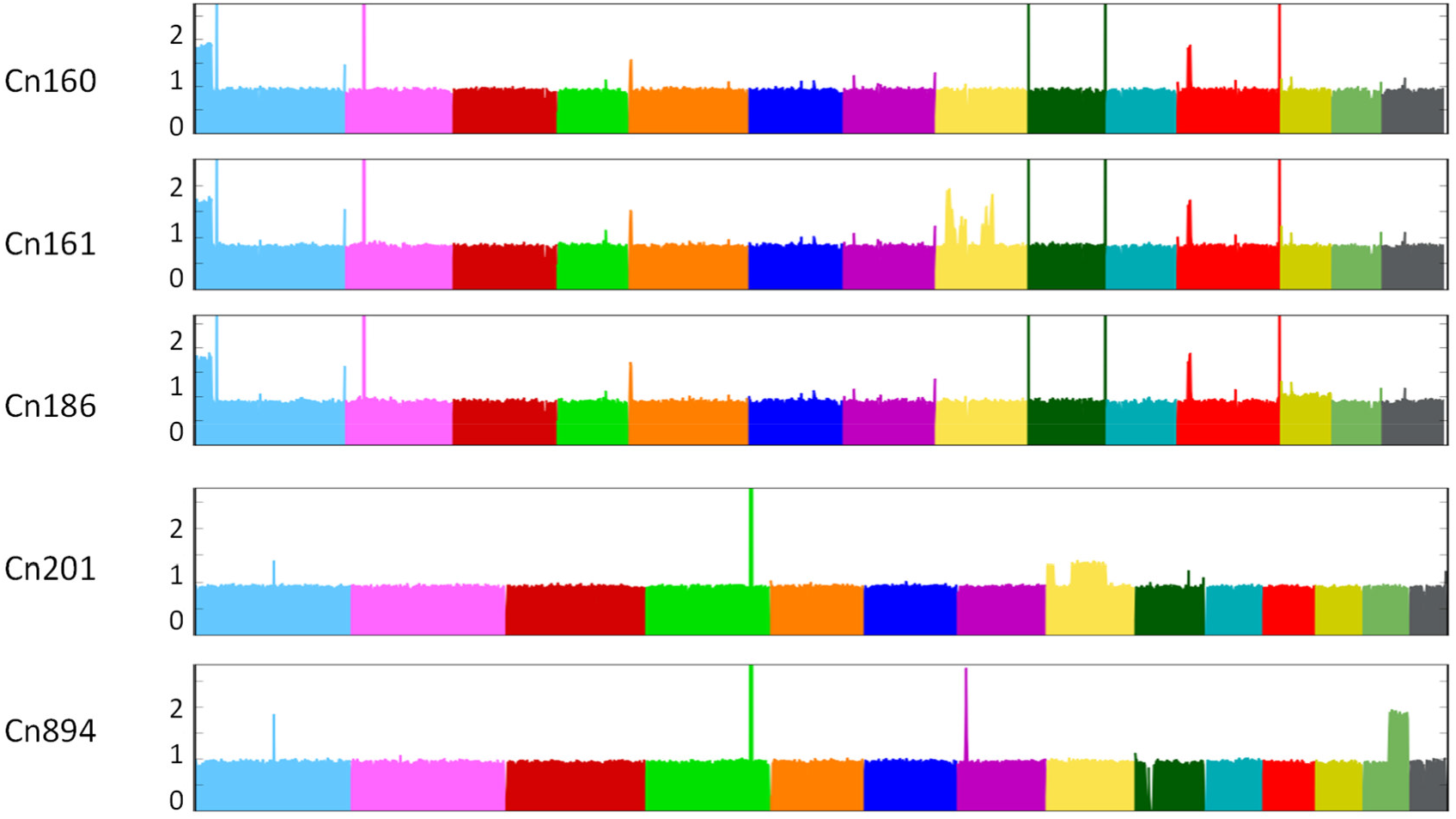
Large segmental duplications within the genomes of Brazilian isolates were revealed after aligning sequencing reads to the *Cryptococcus* reference genomes.

### Phylogenetic analyses

To obtain insight into the population structure of the Brazilian *C. neoformans* isolates, we constructed a phylogenetic tree including 352 isolates considered in previous studies (Desjardins *et al*. 2017; Rhodes *et al*. 2017b; Ashton *et al*. 2019) that encompass all VNI and VNB subclades (Table S2). We also included 10 VNII isolates as the outgroup to root the tree (Desjardins *et al*. 2017; Rhodes *et al*. 2017b; Ashton *et al*. 2019). Overall, 945,565 variable positions were used to construct the tree presented in Figure 2A.

**Figure 2.**
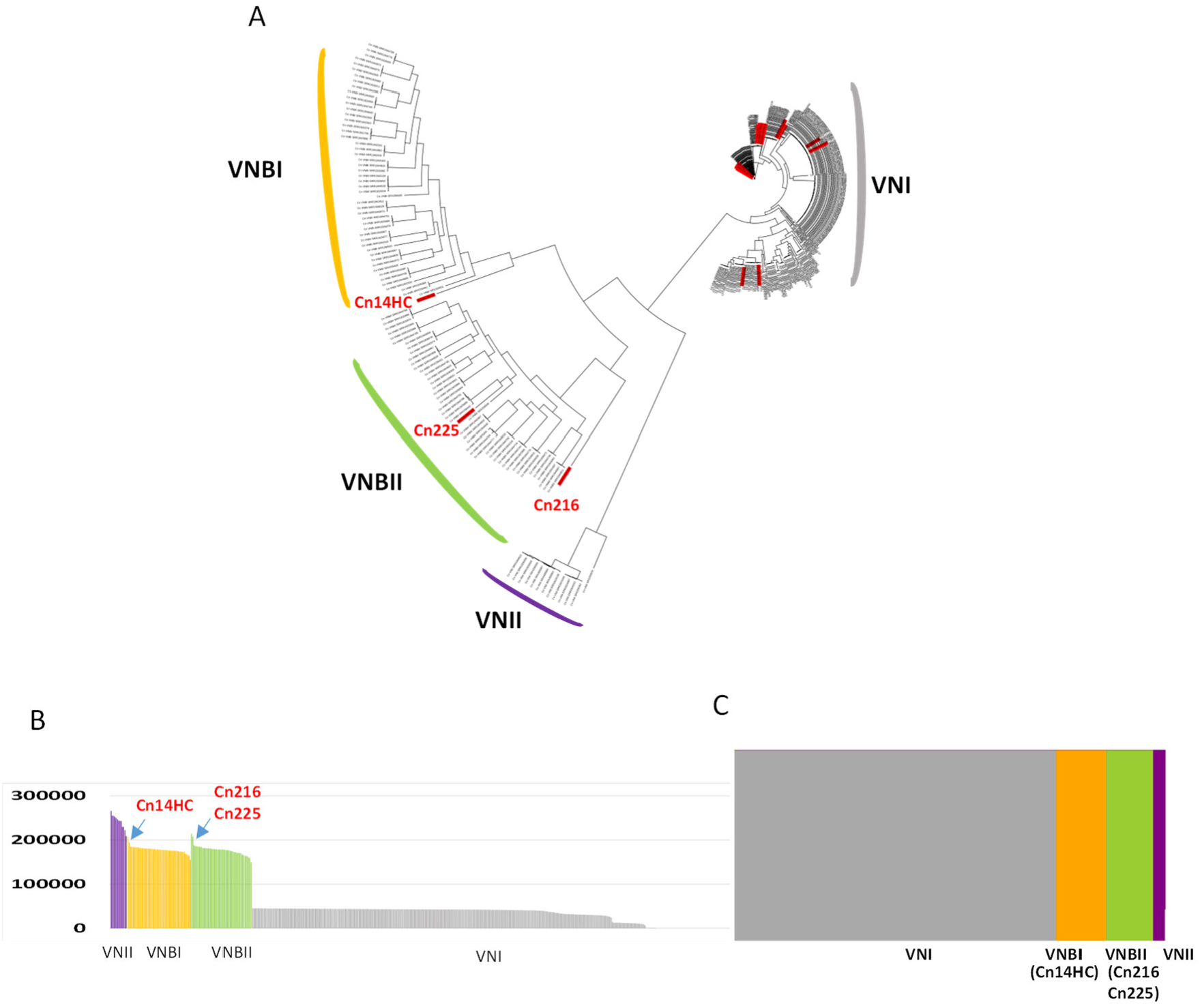
(A) Phylogenetic tree generated by iTOL v5 using *C. neoformans* genome sequences. The Brazilian isolates are indicated in red. The two Brazilian outlier isolates are shown. (B) Number of SNPs within the genomes of the different isolates analyzed in this study. (C) Results of the structure analysis (k=4) suggesting VNBI and VNBII origin for the two Brazilian outlier isolates (CnH14C and Cn216, respectively).

#### VNB isolates

Whereas the vast majority (92.5%, n=37) of the *C. neoformans* Brazilian isolates belonged to the VNI clade, we also identified one VNB isolate of the VNBII subclade (Cn225). Two other Brazilian isolates (Cn216 and Cn14HC) appear to be associated with the VNBI and VNBII clades, respectively, but with a close-to-the-root long branch (Figure 2A). Accordingly, the number of SNPs compared to the H99 reference genome was slightly higher in these two isolates than in the other VNB isolates (Figure 2B). As previously reported, analysis of the population using *fastSTRUCTURE* (k=4) (Raj *et al*. 2014) readily separated VNI, VNII, VNBI, and VNBII isolates (Desjardins *et al*. 2017) into four independent populations (Figure 2C). As expected, this analysis showed that the two Brazilian outlier isolates (Cn216 and Cn14HC) are associated with the VNBI and VNBII clades, respectively.

Because VNI/VNB and VNII/VNB hybrids have been previously reported within African isolates (Rhodes *et al*. 2017b), we sought to compare their phylogenetic position with the two outlier Brazilian isolates. We first redrew a phylogenetic tree including six of these isolates (the VNII/VNB isolate MW-RSA852 and four VNI/VNB isolates [Bt125, Bt131, Bt162, and Bt163]) (Figure S1). As previously published, the African hybrid isolates are also positioned at the root of the VNB and VNII branches, respectively. We then performed principal component analysis (PCA) including this set of hybrid isolates. As previously published, this analysis readily separated isolates positioned at an intermediary position between the defined clades (Figure 3) (Desjardins *et al*. 2017). Like most African VNI/VNB hybrids, the Bt163 isolate was positioned between VNI and VNBII, suggesting that it is a hybrid of isolates of these two clades. Of note, the VNI/VNB isolate Bt125 has an intermediary position between the VNBI, VNBII, and VNI lineages (Figure 3).

**Figure 3.**
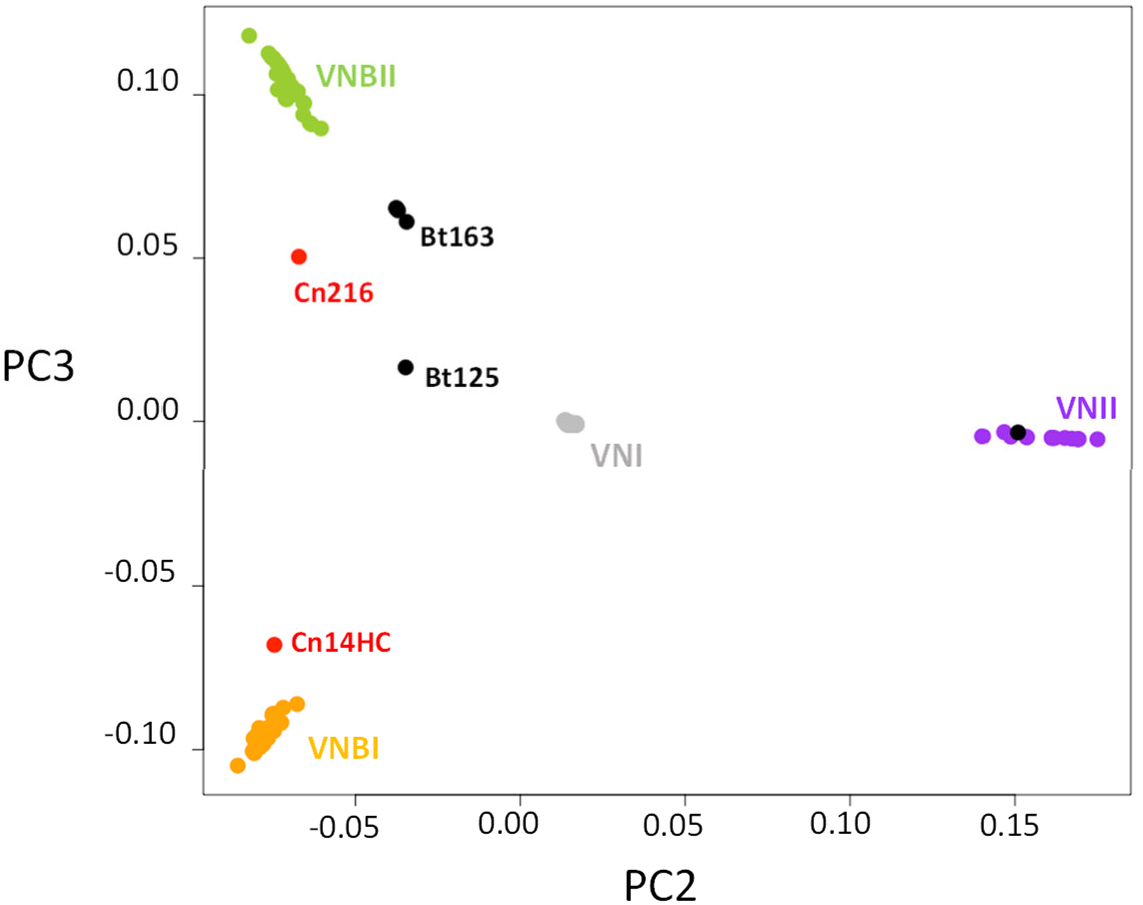
Principal component analysis revealing the intermediary position of the Brazilian outlier isolates. VNI/VNB and VNII/VNB hybrid isolates (Rhodes *et al*. 2017b) have been added to this analysis. The positions of the hybrid (black) and of the two Brazilian outlier isolates (in red) are indicated.

The two outlier Brazilian isolates are positioned between the VNBI and VNBII isolates, suggesting that they are not VNB/VNI hybrids. In agreement with the location of these isolates in the phylogenetic tree, this analysis showed that the Cn216 isolate is closer to VNBII isolates whereas the Cn14HC isolate is closer to VNBI isolates. According to these results, Cn216 and CnH14C could be either VNBI/VNBII hybrids or evolved VNBI and VNBII isolates, respectively. In order to test these two hypotheses, we first determined the position and number of SNPs specific to each clade (Figure 4A,B). This analysis revealed that the number of clade-specific SNPs was equivalent for each clade with an even distribution along the H99 reference genome. Of note, the VNBI clade appear to have fewer clade-specific SNPs, which might be related to the previously reported loss of diversity in this lineage (Desjardins *et al*. 2017).

**Figure 4.**
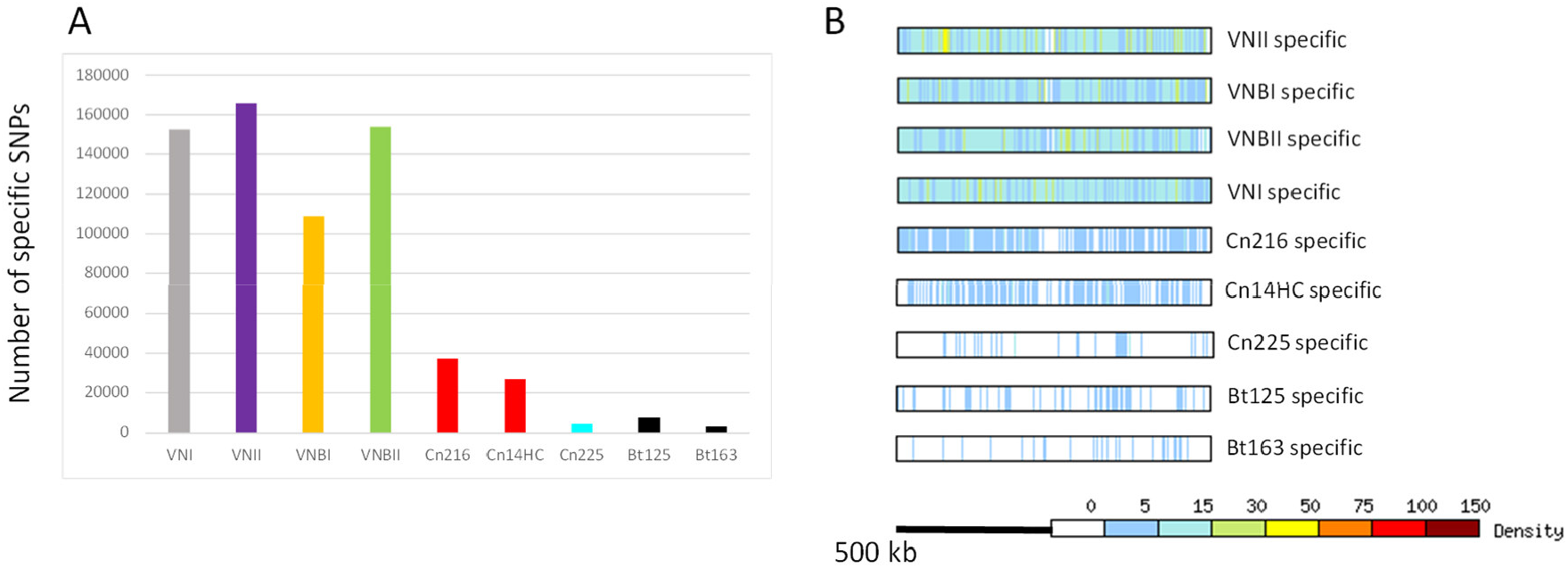
(A) Number of specific SNPs in each clade and in the two Brazilian outlier isolates (Cn216 and Cn14HC). The Brazilian VNBII isolate Cn225 and the hybrid isolates Bt125 and Bt163 (Rhodes *et al*. 2017b) were used as controls. (B) Density and position of the clade-specific SNPs in the H99 reference genome. The results obtained with chromosome 14 are presented.

We reasoned that we could determine the origins of the SNPs in the two outlier Brazilian isolates by comparing them with clade-specific SNPs (Figure 5). We compared the SNPs of Cn216 and Cn14HC SNPs (n=213,915 and n=207,438, respectively) with the VNI-, VNII-, VNBI-, and VNBI-specific SNPs and plotted their positions in the H99 reference genome. As a control, we performed the same analysis with the VNBII Brazilian isolate Cn215 (n=207,639 SNPs).

**Figure 5.**
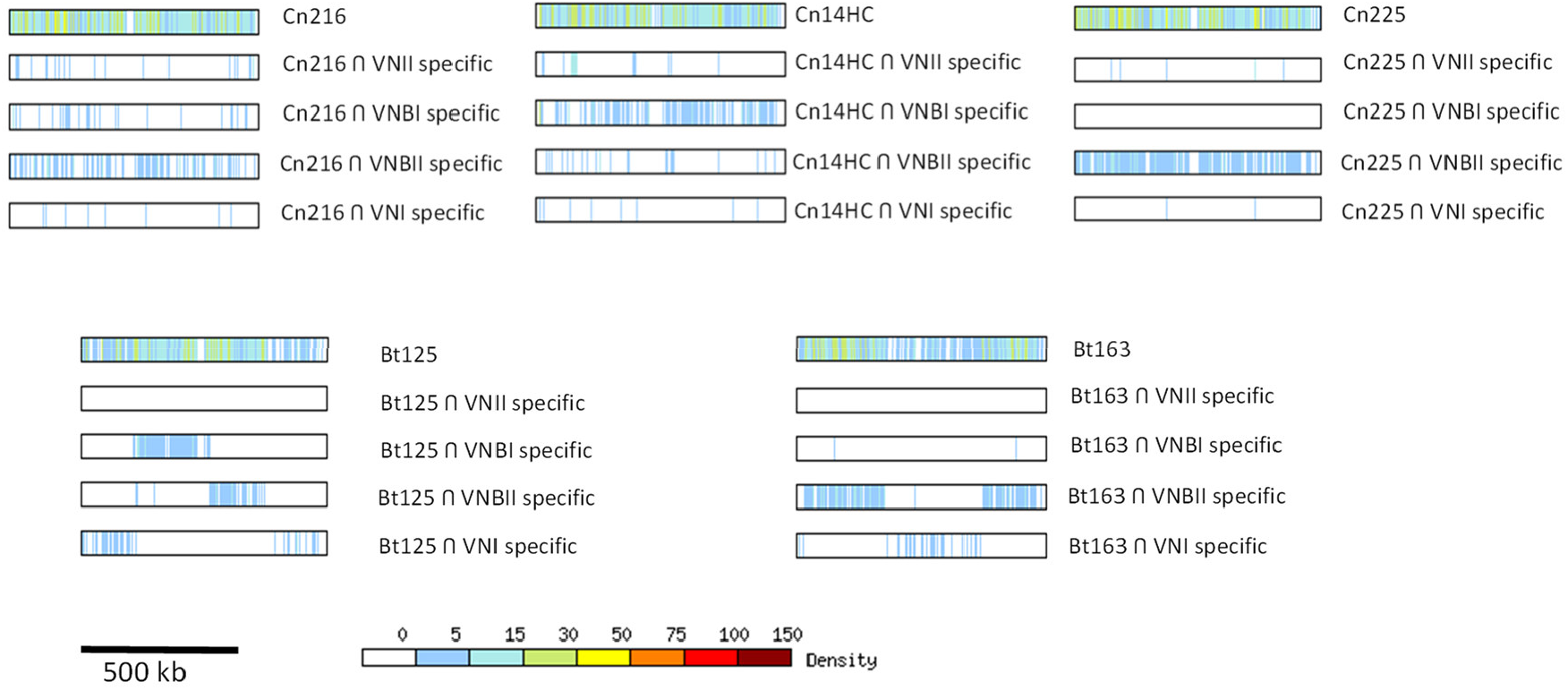
Origin of the SNPs in the Brazilian outlier isolates (Cn216 and Cn14HC). Positions of the SNPs shared between the clade-specific SNPs were identified and plotted on each chromosome. Chromosome 14 is presented as an example. Results obtained with the Brazilian VNBII Cn225 isolate and the African VNB/VNI hybrid isolates Bt125 and Bt163 (Rhodes *et al*. 2017b) are shown as controls.

This analysis revealed that the Brazilian outlier isolates Cn214 and Cn14HC mostly share SNPs with the VNBII- and VNBI-specific SNPs, respectively, confirming that they belong to these respective lineages. We observed an even distribution of these SNPs along the chromosomes with no sign of a recent recombination event between the VNBI and VNBII lineages. As expected, a similar result was obtained with the Brazilian VNBII isolate Cn225. In contrast, the African VNB/VNI hybrid Bt163 displays a SNP distribution common to the clade-specific SNPs, supporting a recent recombination event between a VNBII isolate and a VNI isolate and thus confirming the PCA results and previously published results (Rhodes *et al*. 2017b). For the Bt125 isolate, the SNP pattern was more complex than previously published, suggesting a triple VNBI/VNBII/VNI origin for this isolate, in agreement with the intermediary position of this isolate observed by PCA (Figure 3). Overall, these analyses suggest that the Cn214 and CnH14C isolates belong to the VNBII and VNBI clades, respectively, with no sign of recombination with another clade. Yet, the number of the SNPs unique to these two isolates (n=37,236 in Cn216 and n=26,925 in CnH14C) is much higher than in the VNBII Brazilian isolate Cn225 (n=4788) and in the two VNI/VNB African hybrids (n=7485 in Bt125 and n=3511 in Bt163) (Figure 4A). These isolate-specific SNPs are evenly distributed along the H99 reference genome (Figure 4B). This SNP pattern may be due to an ancient separation of these isolates from most other isolates of their respective clades or a recent but rapid evolution of their genome sequences due to an increased mutation rate.

In order to test these hypotheses, we compared the mutation rates in both Cn216 and Cn14HC by observing the appearance of spontaneous 5-fluorouracil (FU)-resistant colonies in fluctuation assays (Boyce *et al*. 2017; Billmyre *et al*. 2020). We included the reference strain KN99α and the Brazilian VNBII isolate Cn225 as controls. The mutation rate of Cn14HC was higher than those of Cn225 and KN99α (Figure 6). In contrast, the mutation rate in Cn216 was not significantly different from that of Cn225 (VNBII). Similar results were obtained when 5-FOA selection was used (Figure S2).

**Figure 6.**
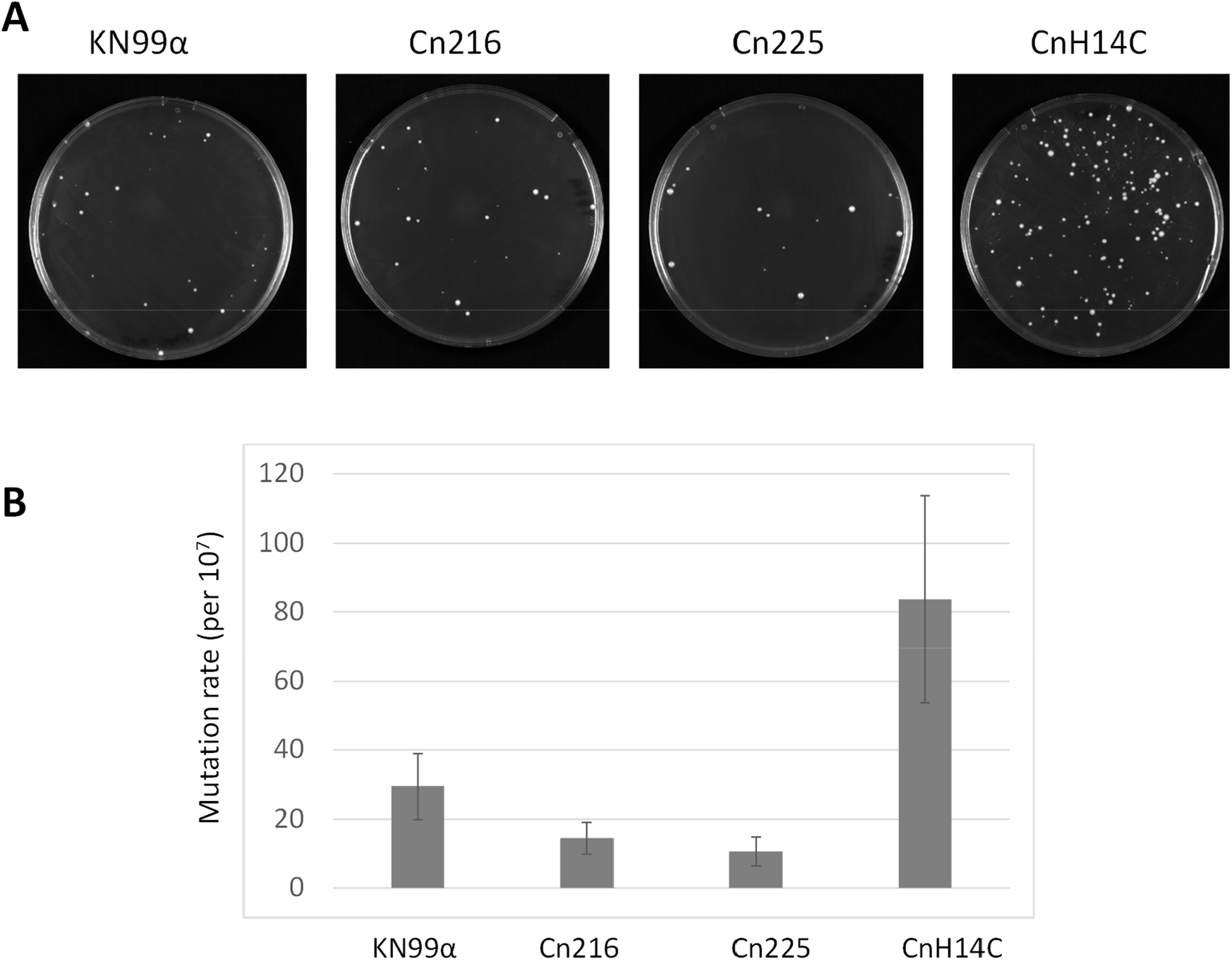
CnH14C display an elevated mutation rate. (A) Spontaneous 5-FU–resistant colonies in a standard wild type (KN99α) isolate and three Brazilian isolates (Cn216, Cn225, and CnH14C). (B) Quantitative assessment of mutation rates in the four strains using fluctuation analysis of the spontaneous resistance to 5-FU. The Lea-Coulson method of the median was used to estimate the number of mutations from the observed values of mutants from 10 independent, parallel cultures (Foster 2006; Gillet-Markowska *et al*. 2015). Error bars indicate 95% confidence intervals.

Isolates with mutations in the mismatch repair pathway associated with a mutator phenotype have been observed both in *C. deuterogattii* (Billmyre *et al*. 2017) and *C. neoformans* clinical isolates (Boyce *et al*. 2017; Rhodes *et al*. 2017a). Although experimental in vitro analyses suggested that only *MSH2, MLH1*, and *PMS1* play a major role in regulating the mutation rate in *C. neoformans* (Boyce *et al*. 2017), recurrent isolate evolution analysis suggested that *MSH5* and *RAD5* could also influence isolate evolution in patients (Rhodes *et al*. 2017a). Given that we did not identify Cn14HC nonsense mutations, we looked at the Cn14HC-specific SNPs responsible for non-conservative mutations in these mismatch repair genes. Interestingly, two of these genes contain mutations specific to Cn14HC: *PMS1* (CNAG_07955) encodes a protein with a S483P mutation and *RAD5* (CNAG_04733) encodes a protein with a P332S mutation. These SNPs were not present in any of the other 391 isolates considered in this study. In contrast, we did not identify specific SNPs responsible for a change in any of these proteins in Cn216 or Cn225.

Overall, our results suggest that CnH14C possesses a mutator phenotype associated with specific mutations in two mismatch repair genes. Of course, more experiments are needed to formally establish a causal relationship between this genotype and this phenotype. Nevertheless, the mutator phenotype of CnH14C has resulted in its rapid evolution, and could explain its position in the phylogenic tree. In contrast, Cn216 might be the result of a long evolution independent of the African VNBII isolate due to an ancient geographical separation.

#### VNI isolates

As stated above, nearly all *C. neoformans* isolates in this collection belong to the VNI clade, mostly within the VNIa subclade. Only two isolates were identified as VNIb and none were identified as VNIc. Interestingly, the second major lineage within the VNIa Brazilian isolates was VNIa_Y, encompassing 27.5% (n=10) of isolates. All of these strains were isolated from pigeon droppings in Northern Brazil. These are the first VNIa_Y strains isolated outside of Africa; previous strains, all of which originated in the environment, were isolated in Botswana, Malawi, and Uganda (Ashton *et al*. 2019). The population structure of the Brazilian VNIa isolates resembles that of the African isolates. Approximately 45% of the VNIa isolates are of the VNIa_93 lineage, similar to previous observations in Ugandan isolates (Ashton *et al*. 2019). Similar to previous reports for African countries, only 2 of the Brazilian isolates were of the VNIa_5 subclass and no VNIa_4 isolates were identified, confirming that these two subclasses are specifically enriched within Southeast Asia isolates (Ashton *et al*. 2019).

Phylogenetic trees constructed using only the VNIa subclades revealed a clear geographical clustering of the isolates (Figure 7), suggesting that the current subclade organization of *C. neoformans* isolates predates the dissemination into South America. However, the existence of unclustered VNI-93 Brazilian isolates suggests multiple intercontinental isolate transfers.

**Figure 7.**
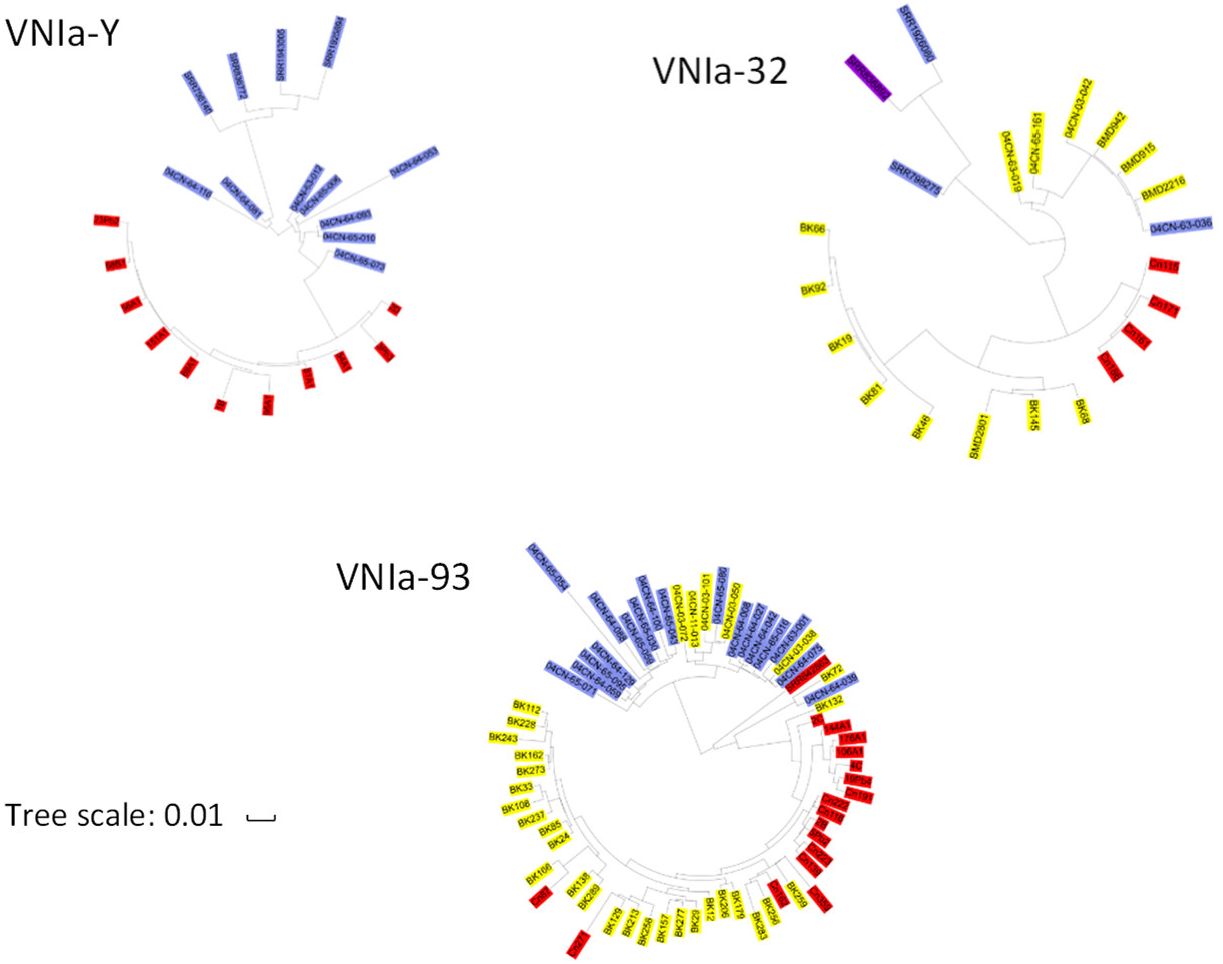
Phylogenetic trees of the VNIa subclades as visualized by ITOL revealed geographic clustering. The Brazilian, African, and South Asian isolates are labelled in red, blue, and yellow, respectively. The Indian isolates are labelled in purple.

#### Phenotypic analyses

We next characterized the Brazilian isolates at a phenotypic level. We focused our analysis on virulence-associated phenotypes and drug resistance. All isolates grew well at 30°C, and only *C. gattii* isolate Cg187 and *C. neoformans* VNIa_Y isolate 3Pb3 exhibited a growth defect at 37°C (Figure S3). Similarly, most of these *Cryptococcus* isolates demonstrated a low MIC for fluconazole (Table S3), with a MIC_50_ of 2 μg/mL and MIC_90_ of 8 μg/mL. The clinical isolate Cn191 (VNIa_93) and the environmental isolate 151A1 (VNIa_Y) showed the highest MIC values (8.0 μg/mL).

Capsule formation and exportation of GXM, a major capsular antigen, are essential for *Cryptococcus* virulence. We analyzed multiple aspects of capsule formation in Brazilian isolates, including capsular morphology, serological reactivity, and extracellular concentration of GXM. Capsule morphology was examined by fluorescence microscopy after incubation of the cells with an antibody to GXM for capsule visualization and calcofluor white for cell wall staining. Alternatively, the capsule was counterstained with India ink for visualization of the capsule by light microscopy. This analysis revealed formidable heterogeneity at morphological and serological levels (Figure 8).

**Figure 8.**
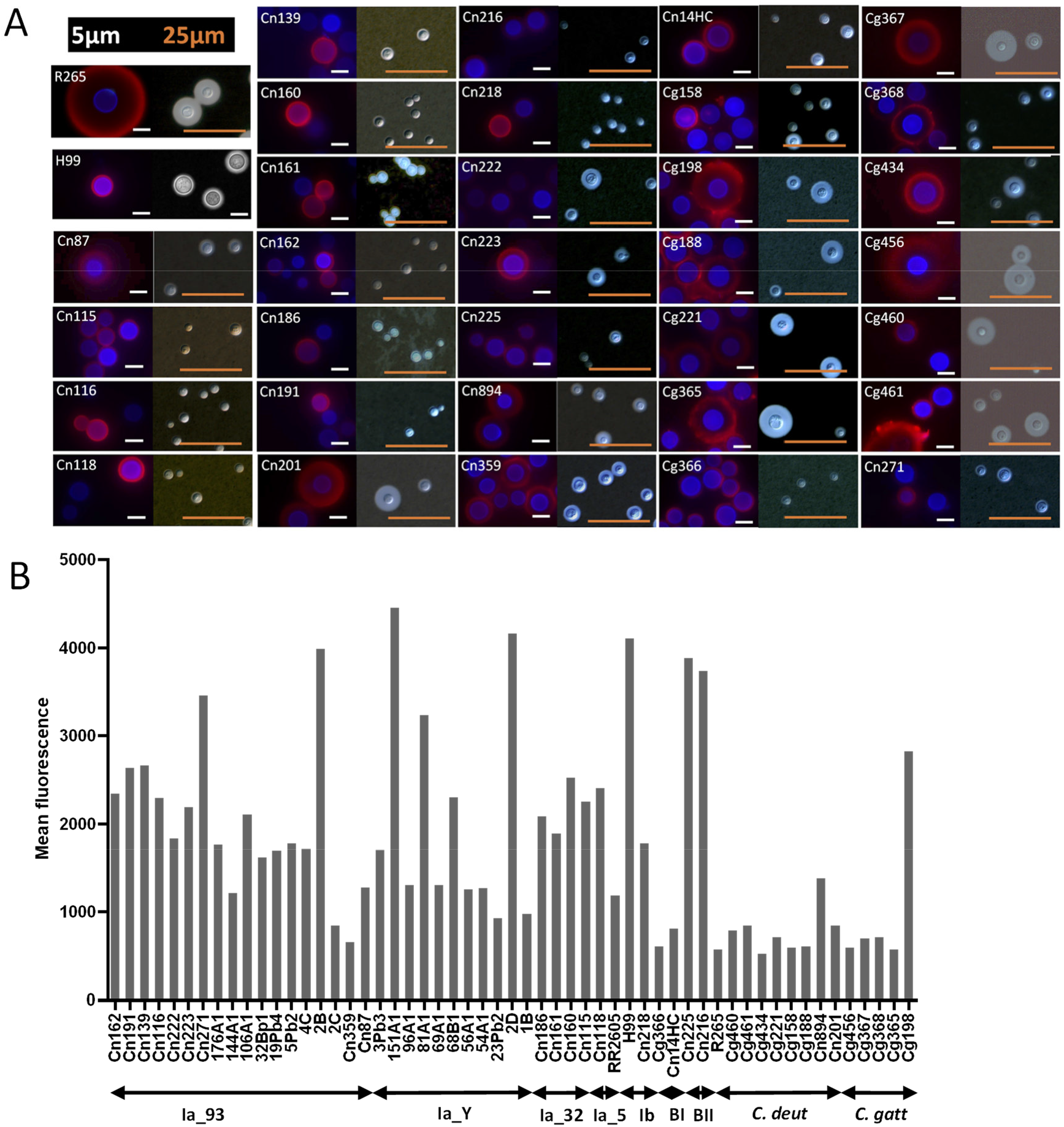
(A) Microscopic analysis of the cell surface of the Brazilian isolates. R265 and H99 are standard cryptococcal isolates that were used as reference in this assay. For each isolate, fluorescence panels are shown on the left and India ink counterstaining is shown on the right. Red fluorescence indicates capsule staining with the mAb 18B7 antibody and blue fluorescence represents cell wall staining. Images are illustrative of two independent analyses that produced similar results. Corresponding data for the Northern Brazil isolates have been previously published (Dal Pupo *et al*. 2019). (B) Flow cytometry analysis of the cryptococcal isolates after capsule staining with mAb 18B7. At least 5000 cells were analyzed for each isolate. Results are presented as average values of fluorescence intensity and are illustrative of two independent analyses producing similar results. R265 and H99 are standard cryptococcal isolates that were used as reference in this assay. Data for the Northern Brazil isolates were obtained from (Dal Pupo *et al*. 2019).

On the basis of the visual perception that antibody recognition of the capsule was variable, we measured the indices of fluorescence intensity after incubation of each isolate with the GXM-binding antibody by flow cytometry (Figure 8B). Using the standard isolates H99 and R265 as reference strains, this analysis confirmed both highly reactive isolates and poorly reactive isolates, supporting the notion that surface GXM is highly diverse in *Cryptococcus*. Although *C. gattii* and *C. deuterogattii* isolates were predominantly less capsulated than *C. neoformans* (student t-test *P* value <0.05), we did not observe any statistical difference between the *C. neoformans* subclades.

We also analyzed the concentration of polysaccharide in culture supernatants of each isolate (Figure S4). Detection of extracellular and surface-associated GXM showed an apparent correlation, as suggested by the statistical analysis (Figure S4), supporting the notion that cryptococcal isolates are highly variable in their ability to form GXM-containing capsules and to export polysaccharide.

We also examined melanin production on L-Dopa medium. We observed a variable phenotype among the Brazilian isolates, with *C. gattii* and *C. deuterogattii* isolates displaying lower melanin production (Figure 9). There was no obvious correlation between melanin production among subclades, although strains isolated from bird droppings appear to produce more melanin than other environmental or clinical isolates.

**Figure 9.**
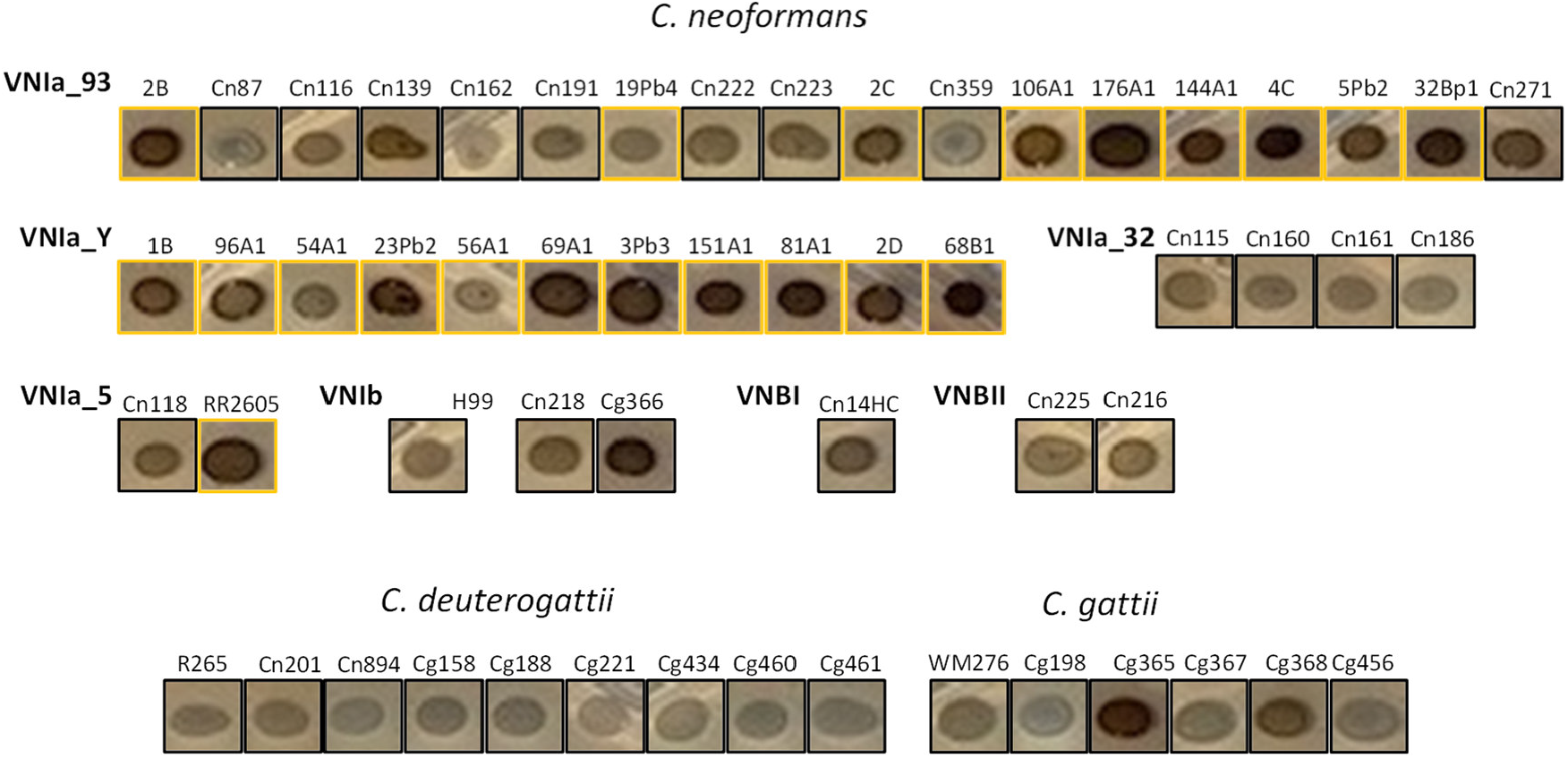
Melanin production of Brazilian isolates as estimated on L-Dopa medium after 72 h of incubation. Strains isolated from bird droppings are indicated by a yellow box.

Lastly, we examined production of TCs. *C. neoformans* and *C. gattii* cells can undergo an unusual transition in the host lung from the typical size of 5 to 7 μm to between 10 μm and 100 μm, a phenomenon known as titanization. TCs are known to be highly polyploid (Zaragoza and Nielsen 2013). TCs can also be induced in vitro using three different approaches (Dambuza *et al*. 2018; Hommel *et al*. 2018; Trevijano-Contador *et al*. 2018). Here, we followed the protocol described by Dambuza and colleagues (2018) to study whether Brazilian *Cryptococcus* isolates produce TCs. In our study, *C. neoformans* H99 formed nearly 17% TCs, aligning with previously published findings (Dambuza *et al*. 2018). Approximately 49% of Brazilian *Cryptococcus* isolates produced TCs (defined here as cells with a size >10 μm) (Figure 10, Table S4). Around 24% of Brazilian *Cryptococcus* isolates generated more TCs than *C. neoformans* H99 (Figure 10, Table S4). Interestingly, and in contrast to a previous study using the same protocol, all Brazilian *C. gattii* and *C. deuterogattii* isolates tested formed TCs (Figure 10, Table S4). The highest percentage of TCs (97.33%) was produced by *C. deuterogattii* isolate Cn894 (Figure 10, Table S4).

**Figure 10.**
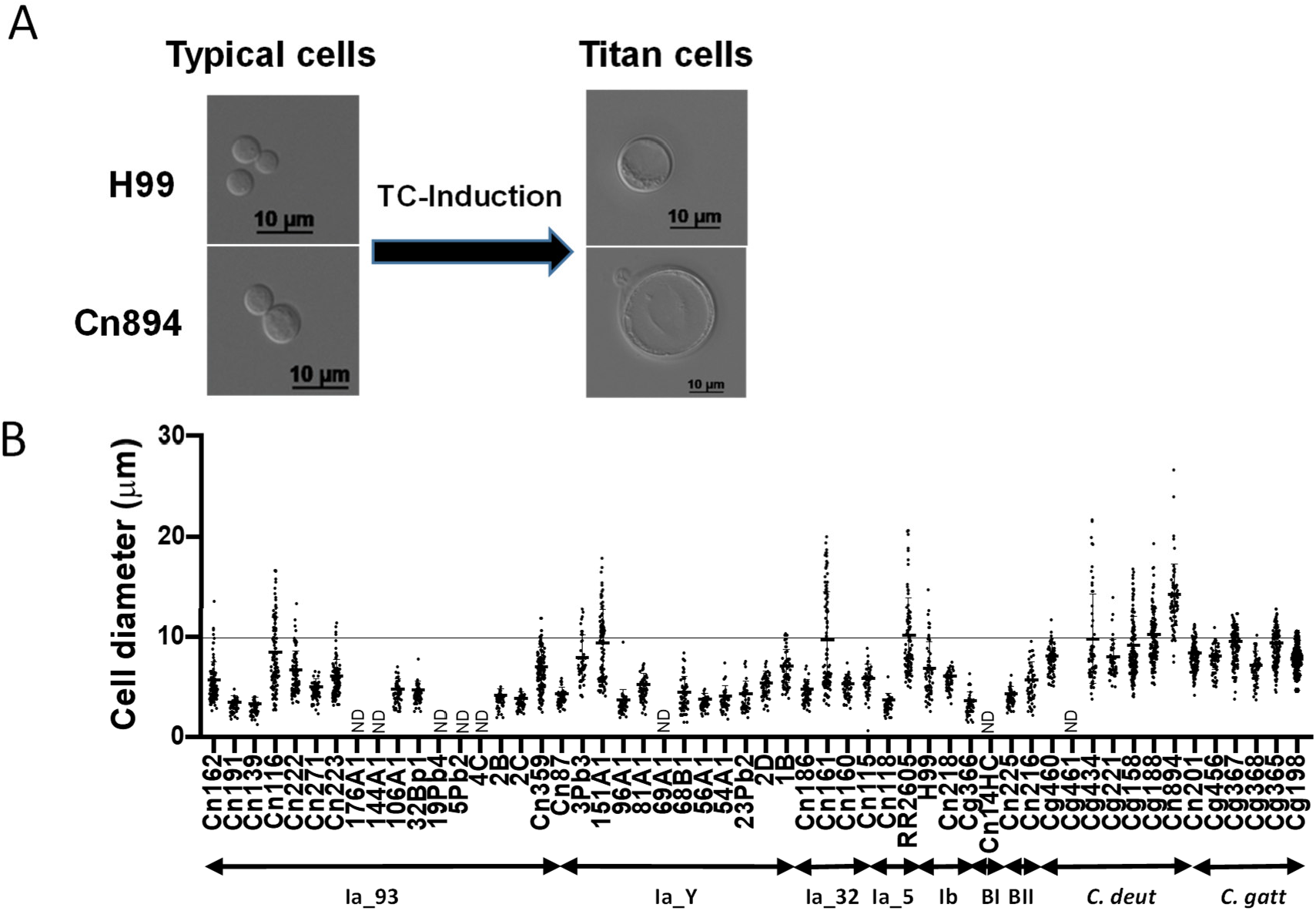
Cell diameter after titanization induction in Brazilian isolates. Cryptococcus isolates were subjected to TC induction in PBS-HI-FCS. Cells with a diameter >10 µm were considered to be TCs. ND: not determined.

In order to address whether cells that formed more TCs than H99 are naturally larger, we compared the size of these cells before and after the titanization induction. The mean size of the cells of Brazilian *Cryptococcus* isolates varied between 4.18 to 5.70 μm before and 6.85 to 14.19 μm after cultivation in TC-inducing conditions, proving that the titanization induction was responsible for the increase in cell size (Table S5). We did not observe any link between subclade and the ability of the corresponding isolate to undergo this phenotypic change.

Overall, this analysis revealed spectacular phenotypic variability within the Brazilian isolates, although no association was detected between one clade and any of these phenotypic traits.

## Discussion

Population genomics studies have resulted in a model in which clonal reproduction has shaped the global population structure of *C. neoformans* isolates, which are mostly of the VNI clade and of the *MATα* sexual type (Cuomo *et al*. 2018). These genome sequence analyses have also identified several VNI subclades with specific geographical distribution (Ashton *et al*. 2019). Currently, little is known about *Cryptococcus* species diversity in South America because only a few isolates have been sequenced (Rhodes *et al*. 2017b). In the present study, we analyzed the largest South American genome collection studied to date. As expected, most of the Brazilian isolates belonged to the VNI clade. Our analysis revealed that the VNI Brazilian *C. neoformans* population shares many features with the African population. For instance, VNI_Y isolates that were previously reported only in Africa were also prominent within this set of South American isolates. As in Africa, Brazilian VNIa_5 isolates were rare; in contrast, VNIa_5 isolates represent one of the largest subclades within Asian isolates (Ashton *et al*. 2019). Actually, in most regards, the *C. neoformans* Brazilian population structure is very different from the Asian one described by Ashton and colleagues in 2019 (Ashton *et al*. 2019). A detailed analysis of the different VNIa subclade population structures further supported the hypothesis of repeated and privileged exchanges from African to South American isolates. Thus, South American isolates do not form an independent clade or subclade but rather are scattered within nearly all global subclades, indicating repeated transport. Although we observed a perfect clustering of Brazilian isolates within the VNIa_Y and VNIa_32 subclades—possibly from an ancient importation followed by independent evolution of the ancestral isolate—some VNIa-93 isolates clustered with the other global isolates, suggesting independent and possibly more recent importations. Of course, because, all these unclustered VNI-93 Brazilian strains are clinical isolates, we cannot completely exclude the hypothesis that these patients had travelled to Africa, and had been infected there, before developing a cryptococcosis in Brazil as previously described in some rare cases (Garcia-Hermoso *et al*. 1999).

A similar pattern was observed with the VNB isolates. Two of the Brazilian isolates were positioned close to the roots of the VNB branches, as previously observed for two sequenced Brazilian isolates (Rhodes *et al*. 2017b). However, the third isolate clustered within the other VNB African isolates, suggesting a more recent intercontinental exchange. In that case, it is important to note that this last strain has been isolated in the environment.

In 2017, Casadevall and colleagues suggested that the separation of *C. neoformans* complex species and *C. gattii* complex species occurred in the middle of the Cretaceous period (Casadevall *et al*. 2017). This geological period was marked by the breakup of the supercontinent west Gondawa which encompassed Africa and South America. This modern continent formation ended much before the global dispersion of major *Cryptococcus* spreader birds (Prum *et al*. 2015) which occurred 40 × 10^6^ years after the end of the Cretaceous period. Although our data cannot reliably date the separation between the *C. neoformans* subclades, the similar structure of the African and South American population suggests an African origin for every *C. neoformans* isolates. In this model, the transport of *C. neoformans* from Africa to South America started only with birds migrations 25 × 10^6^ years ago and continued until today. This study also exemplified the impact of rapidly evolving isolates on the *C. neoformans* population structure. Mutator phenotypes have already been observed in natural *C. neoformans* and *C. gattii* isolates and have been shown to impact the dynamics of isolate evolution, phenotypic diversity, and drug resistance (Billmyre *et al*. 2017; Billmyre *et al*. 2020). A *msh2*-associated mutator phenotype was also observed in a subset of *C. deuterogattii* Pacific Northwest outbreak isolates, named VGIIa-like (Kidd *et al*. 2004; Billmyre *et al*. 2014; Billmyre *et al*. 2017). Here, the VNBI isolate CnH14C displayed a mutator phenotype, albeit less pronounced than the previously described *C. neoformans msh2Δ, mlh1Δ*, and *pms1Δ* mutant strains (Boyce *et al*. 2017). Accordingly, we did not identify nonsense mutations in any of these genes in the CnH14C isolate. However, *PMS1-* and *RAD5*-specific alleles in CnH14C might suggest a causal relationship. In fact, *RAD5* has been recently identified as regulating mutation rates in a quantitative trait locus analysis in *S. cerevisiae* (Gou *et al*. 2019), and a specific *RAD5* mutation has also been identified in a rapidly evolved *C. neoformans* recurrent clinical isolate (Rhodes *et al*. 2017a).

However, in contrast to what has been observed in some African isolates, we did not identify any hybrid isolates within this collection of Brazilian isolates. As previously reported (Matsumoto *et al*. 2007; Andrade-Silva *et al*. 2018), none of these isolates were *MAT***a**. Nevertheless, our model of multiple intercontinental transport of African isolates to South America and the proportion of *MAT***a** isolate within the VNB isolates in Africa (Litvintseva *et al*. 2007; Litvintseva *et al*. 2011; Litvintseva and Mitchell 2012) predict that additional sampling will probably identify such isolates.

MLST and genome sequence analysis identified South America, and in particular Brazil, as the major source of the global genetic diversity of the *C. gattii* complex, especially within the VGII and VGIII lineages (Hagen *et al*. 2013; Billmyre *et al*. 2014; Souto *et al*. 2016; Barcellos *et al*. 2018; Vilas-Bôas *et al*. 2020). This is associated with great phenotypic diversity within the Brazilian isolates (Silva *et al*. 2012; Barcellos *et al*. 2018). Similarly, genotypic and phenotypic analyses of Brazilian *C. neoformans* isolates, including the analysis in the present study, have revealed great diversity within South American isolates (Ferreira-Paim *et al*. 2017; Andrade-Silva *et al*. 2018; Dal Pupo *et al*. 2019; De Sousa *et al*. 2020). Nevertheless, our data suggest that Brazil is probably not an important source of global diversity of *C. neoformans* isolates. Rather, our findings support the model in which this diversity is mostly imported from Africa through repeated intercontinental transport of *C. neoformans* isolates to South America. Our studies continue the investigation of *Cryptococcus* species global diversity and emphasize the importance of understanding the evolution of fungal pathogens. Further studies with additional isolates from Brazil and other countries in South America will help us to clarify many aspects of population genetics of this important fungal pathogen.

## Acknowledgments

M.L.R. is currently on leave from the position of associate professor at the Microbiology Institute of the Federal University of Rio de Janeiro, Brazil. M.L.R., G.J., and G.H.G. were supported by a Tripartite USP-Fiocruz-Pasteur grant. FCGR was financed in part by a scholarship from the Coordenação de Aperfeiçoamento de Pessoal de Nível Superior (CAPES, Brazil, Finance Code 001). This work in the Janbon lab was supported in part by an Infect-ERA grant (project Cryptoview).

